# X-linked meiotic drive boosts population size and persistence

**DOI:** 10.1101/2020.06.05.137224

**Authors:** Carl Mackintosh, Andrew Pomiankowski, Michael F Scott

## Abstract

X-linked meiotic drivers cause X-bearing sperm to be produced in excess by male carriers, leading to female-biased sex ratios. Selection for these selfish sex chromosomes can lead to completely female populations, which cannot produce offspring and go extinct. However, at the population level, moderately female-biased sex ratios are optimal because relatively few males are required to fertilise all the females. We develop eco-evolutionary models for sex-linked meiotic drive alleles to investigate their full range of demographic effects. We find general conditions for the spread and fixation of X-drivers, accounting for transmission bias and other factors associated with the spread of X-drivers such as sperm competition and polyandry. Our results suggest driving X-alleles that do not reach fixation (or do not bias segregation excessively) will boost population sizes and persistence times by increasing population productivity, demonstrating the potential for selfish genetic elements to move sex ratios closer to the population-level optimum. We suggest that researchers should look beyond extinction risk and consider the potential for ecologically beneficial side effects of selfish genetic elements, especially in light of proposals to use meiotic drive for biological control.

## Introduction

Meiotic drivers violate Mendel’s law of equal segregation by ensuring that they are transmitted to more than half of a carrier’s progeny (Burt and Trivers 2006). While beneficial at the chromosome-level, this transmission benefit usually comes at a cost to their carrier’s survival or fecundity (Werren 2011). Meiotic drive has been observed across a wide variety of animal and plant taxa (Sandler *et al.* 1959; Turner and Perkins 1979; Jaenike 1996; Ardlie 1998; Taylor *et al.* 1999; Fishman and Willis 2005; Tao *et al.* 2007; Lindholm *et al.* 2016), particularly in flies and rodents (Helleu *et al.* 2015). Many of the described systems are sex-specific (Úbeda and Haig 2005; Lindholm *et al.* 2016), arising due to drivers that are active during meiosis in either females (e.g., Fishman and Willis (2005)) or males (e.g., Sandler *et al.* (1959)). When meiotic drivers arise on sex chromosomes, they change the relative frequencies of gametes carrying the sex-determining alleles, which causes the sex ratio at birth to become biased (Burt and Trivers 2006). In particular, where X-linked meiotic drivers bias segregation in males, X-bearing sperm outnumber Y-bearing sperm and the sex ratio among offspring is female-biased. Hamilton (1967) noted that extreme sex ratios caused by X-linked meiotic drivers could lead to population extinction, as eventually the almost entirely female population will go unmated and be unable to produce offspring.

Substantial theoretical work since Hamilton’s pioneering study (Hamilton 1967) has investigated the spread of meiotic drive, and the conditions that lead to its polymorphism and prevent population extinction. Polymorphism and population persistence are most directly achieved via suppression systems that evolve at other loci to negate meiotic drive (Hamilton 1967; Charlesworth and Hartl 1978; Frank 1991). In the absence of suppression, fixation of autosomal (Ardlie 1998; Larracuente and Presgraves 2012) or X-linked (Taylor and Jaenike 2002, 2003; Price *et al.* 2014) meiotic drive can be prevented by direct fitness costs associated with carrying the driving allele. Meiotic drive systems often occur within inversions that link together the required drive and enhancer loci (Pomiankowski and Hurst 1999). These inversions may also capture deleterious alleles and/or allow deleterious mutations to accumulate through Muller’s ratchet, potentially explaining the fitness costs associated with meiotic drivers (Edwards 1961; Curtsinger and Feldman 1980; Dyer *et al.* 2007; Kirkpatrick 2010). Such effects have been demonstrated empirically, with female carriers of X-linked meiotic drive observed to have reduced survival or fecundity, especially when homozygous (Larner *et al.* 2019; Dyer and Hall 2019; Keais *et al.* 2020). However, these fitness costs are not necessarily sex-specific or recessive (Finnegan *et al.* 2019b).

Meiotic drive can also have deleterious effects by reducing male fertility, most obviously because sperm/spores that do not carry the driving element are rendered dysfunctional or killed (Price *et al.* 2008). This effect may be negligible when females mate with a single male, but drive can alter competition between the ejaculates of different males in a polyandrous mating system. Not only do drive-carrying males deliver fewer sperm per ejaculate, but drive-carrying sperm can also perform more poorly in sperm competition with sperm from wild-type males (Price *et al.* 2008; Manser *et al.* 2017; Dyer and Hall 2019). Offspring sired by drive males have lower fitness which may favour the evolution of increased sperm competition through female polyandry, an argument for which there is some theoretical and experimental evidence (Price *et al.* 2008; Wedell 2013; Price *et al.* 2014; Holman *et al.* 2015; Manser *et al.* 2017), but see (Sutter *et al.* 2019). The fertility cost to drive males, and associated selection for female polyandry, becomes less important as the male frequency declines and they are less likely to be competing for mates and fertilisation (Taylor and Jaenike 2002, 2003). In line with this, modelling has shown that polyandry can limit the spread of meiotic drive alleles, but that evolution of polyandry is not sufficient to stop meiotic drive alleles fixing (Holman *et al.* 2015).

The above models have focused on the evolutionary dynamics of meiotic drive but ignored its demographic consequences. This is surprising as in one of the foundational models of the field, Hamilton (1967) showed that sex-linked drive causes transient population expansion before extinction. Population decline occurs when the sex ratio is pushed beyond the point where females can find sufficient mates. This model did not include density-dependent population regulation or fertility/viability costs associated with meiotic drive. Nevertheless, it suggests that X-linked meiotic drivers will increase population size when they cause sex ratios to be biased, but not extremely biased. Some subsequent analyses support this hypothesis, but it has not been examined directly. Unckless and Clark (2014) showed that species with X-linked meiotic drivers can have an advantage during interspecific competition, shifting the community competition in their favour (James and Jaenike 1990). Similar effects can occur with other systems that cause female-biased sex ratios. For example, feminisation caused by *Wolbachia* can increase population size until females go unmated due to a lack of males (Hatcher *et al.* 1999; Dyson and Hurst 2004). Finally, under temperature-dependent sex determination, shifts in climate can bias the sex ratio towards females (West 2009), which is predicted to increase population sizes providing males are not limiting (Boyle *et al.* 2014).

Incorporating polyandry, sperm competition, and male transmission bias, we develop a general model for determining X-linked allele polymorphism criteria and demographic equilibria. We complement this with a simulation based model which together indicate that the continued presence of drive can lead to larger, more persistent populations. The resulting populations may be able to survive in marginal habitats where they would otherwise become extinct.

## Materials and Methods

We model a well-mixed population with XY sex-determination where generations are discrete and non-overlapping. There are two types of X chromosome segregating in the population, a standard X chromosome and a drive X_*d*_ chromosome. There are three female genotypes *XX*, X_*d*_X and X_*d*_X_*d*_, and two male genotypes XY and X_*d*_Y, which we describe as wild-type and drive males respectively.

Male fertility is dependent on genotype. XY males contribute one unit of sperm per mating, of which half carry an X and half a Y chromosome. The fertility of drive males is given by *c* (*c* ∈ [0, 1]), where *c* = 1 means no fertility loss and *c* < 1 means there is some loss of fertility relative to standard males. We assume ejaculate sizes are large such that males always produce sufficient sperm to fertilise a female. The X_*d*_ chromosome biases segregation such the ratio of X_*d*_ to Y chromosomes among their sperm is (1 + *δ*)/2: (1 − *δ*)/2. When *δ* = 0, meiosis is fair and sex chromosomes are transmitted with equal probability; when *δ* = 1 males produce only X_*d*_ sperm. In the case *c* = 1/(1 + *δ*), drive acts as a “sperm-killer”, decreasing the quantity of Y sperm without compensating for this loss by producing more X_*d*_ sperm.

We track the genotypes of adults, who mate at random before producing offspring. The number of offspring produced is subject to competition between adults (see Supplementary Information for an alternative form of population regulation). Offspring then experience viability selection before they become the adults of the next generation. We assume that a female’s eggs are not fertilised and laid until the mating phase is over, i.e. competition occurs among the ejaculates of all males a female has mated with.

### Analytical model for polymorphism criteria

In this model, we consider various degrees of polyandry determined by a fixed integer *λ_f_*: females mate *λ_f_* times, with a male mate chosen uniformly at random.

We model density-dependent population regulation by assuming that female fecundity declines linearly with population size due to competition for resources. Specifically, female fecundity is given by *B_N_* = *b*(1 − *αN*), where *b* is the intrinsic female fecundity in the absence of competition and *α* is the per-individual competitive effect on fecundity. The total number of adults in the population is given by *N* = ∑_*i*_ *F*_*i*_ + ∑_*j*_ *M*_*j*_, where *F_i_* and *M_j_* represent female and male population densities respectively and *i* ∈ {*XX*, *XX_d_*, *X_d_, X_d_*} and *j* ∈ {*XY*, *X_d_Y*}.

We assume that genotypes affect fitness via the probability of female and male offspring surviving to adulthood, described by 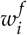 and 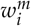. Female and male fitness effects are given by *s_f_*, *s_m_* ∈ [0, 1] and *h* ∈ [0, 1] determines dominance in females (Table 1).

**Table 1.** Relative fitness and transmission parameters for different male and female genotypes

| female genotype $i$ | $w_i^f$ | $E_{X,i}$ | $E_{X_d,i}$ | |
| --- | --- | --- | --- | --- |
| $i = XX$ | 1 | 1 | 0 | |
| $i = X_dX$ | $1 - hs_f$ | 1/2 | 1/2 | |
| $i = X_dX_d$ | $1 - s_f$ | 0 | 1 | |
| male genotype $j$ | $w_j^m$ | $S_{X,j}$ | $S_{X_d,j}$ | $S_{Y,j}$ |
| $j = XY$ | 1 | 1/2 | 0 | 1/2 |
| $j = X_dY$ | $1 - s_m$ | 0 | $c(1 + \delta)/2$ | $c(1 - \delta)/2$ |

When each female mates once (*λ_f_* = 1), the female densities in the next generation are given by

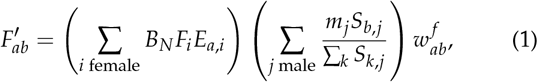

and the male densities by

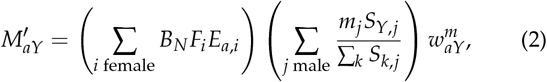

where *m_j_* = *M_j_*/∑_*k*_ *M_k_* is the frequency of males with genotype *j*, *E_a,i_* is the proportion of eggs with haploid genotype *a* produced by females with genotype *i* and *S_b,j_* is the proportion of sperm with haploid genotype *b* contributed by male of X-chromosome genotype *j* (Table 1). The maternal and paternal chromosomes inherited are represented by subscripts *a* and *b*, respectively. As there are no parent-of-origin effects, the sum of 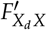 and 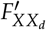 is represented simply as 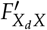.

When each female mates twice (*λ_f_* = 2), female densities in the next generations are given by

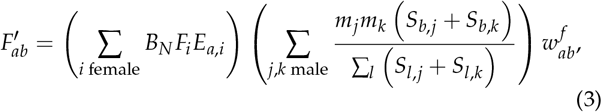

where there is competition for fertilization of each egg among the sperm contributed by two males, firstly with genotype *j* and then with genotype *k*. When each female mates many times (*λ_f_* large), the female densities in the next generation approach

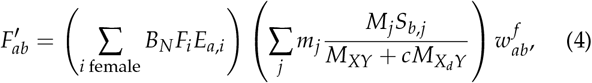

where females effectively sample sperm randomly from the total pool of gametes produced by all males in the population. Recursion equations for male densities follow similarly, replacing *S_b,i_* with *S_Y,i_* and 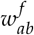with 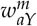 in equations Eq(3) and Eq(4).

### Simulation model

The previous model assumed that male matings are not limiting. Therefore, population extinction could only occur when the birth rate is low and/or no males remain. In this model, we allow limitations on the mating rate in both female and male matings which are capped by *λ_f_* and *λ_m_* respectively. When an individual reaches the maximum number of matings they can no longer mate or be chosen as a mate. So, it is possible for a population to become extinct because the sex ratio becomes female biased and there are insufficient males to sustain the population.

As in the analytical model, female fertility is density-dependent. In the absence of any competition, females lay *b* eggs each. In the case where *b* is non-integer, females lay a mean of *b* eggs by laying a minimum of ⎿*b*⏌ eggs with a 100(*b* − *b*)% chance of laying one more. Whether or not the a birth occurs depends on the competitive influence of other adults, with birth probability 1 − *αN*.

The first generation comprises *N*_0_ wild-type individuals at an equal sex ratio, and the driving X_*d*_ chromosome is introduced into the population at a proportion *q* in Hardy-Weinberg equilibrium. Generations then proceed similarly to the previous model. Adults mate randomly until there are either no females or no males available to mate. Assuming they are able to mate, every individual is picked with equal probability. We track the sperm carried by each female as a 3-tuple (*x*, *y*, *z*), where X, Y, and *z* represent X, X_*d*_, and Y bearing sperm. XY males add (0.5, 0, 0.5) to the pool when they mate, and X_*d*_Y males add (0, *c*(1 + *δ*/2), *c*(1 − *δ*)/2). Once mating is complete, each egg is fertilised by a sperm sampled randomly, weighted by the probability distribution (*x*, *y*, *z*) after normalisation. The juveniles then undergo viability selection, with survival probabilities given in Table 1.

There are three main sources of stochasticity present within the simulation model but not in the analytical model. First, the exact sperm that fertilises an egg is sampled at random. Second, juvenile survival to adulthood and the realisation of births is probabilistic. And finally, mating is at random. These three sources can result in fluctuations in genotype frequencies, which can affect the population sex ratio and population size.

### Data availability

Mathematica notebooks for the main text and supplementary information can be found in Files S1 and S2, and the Python script used to simulate data can be found in File S3 at (figshare link).

## Results

### Maintenance of polymorphism

For a driver locus to remain polymorphic, a driving X_*d*_ chromosome must increase in frequency when rare but not fix in the population. In general, a rare X chromosome allele increases in frequency if

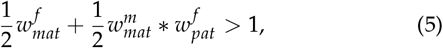

where 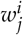 is the relative fitness of the mutant X chromosome in sex i when inherited maternally (*j* = *mat*) or paternally (*j* = *pat*). These relative fitnesses include any transmission biases that arise during gamete production or competition, relative to the transmission of the resident chromosome in the same sex (see Table 3).

**Table 2.** Table of terms

| Term | Definition |
| --- | --- |
| $M_i$ | Density of males with genotype $i$ |
| $F_i$ | Density of females with genotype $i$ |
| $m_i$ | Within-sex frequency of males with genotype $i$ |
| $\lambda_f$ | Number of matings before laying eggs in a females' lifetime |
| $E_{a,i}$ | Proportion of eggs of genotype $a$ produced by a female of genotype $i$ |
| $S_{b,i}$ | Units of sperm of type $b$ produced by a male of genotype $i$ relative to wild-type |
| $N$ | Total density of males and females |
| $\alpha$ | Per-adult cost to average female fecundity |
| $b$ | Intrinsic female fecundity (in the absence of competition) |
| $B_N$ | Female fecundity in a population of size $N$ |
| $c$ | Fertility of a drive male compared to a wild-type male |
| $\delta$ | Strength of drive |

**Table 3.** Values for terms in the general form of the invasion and fixation conditions

| wild-type resident | $\lambda_f = 1$ | $\lambda_f = 2$ | $\lambda_f \text{ large}$ |
| --- | --- | --- | --- |
| $w_{mat}^f$ | $w_{X_d X}$ | $w_{X_d X}$ | $w_{X_d X}$ |
| $w_{mat}^m$ | $w_{X_d Y}$ | $w_{X_d Y}$ | $w_{X_d Y}$ |
| $w_{pat}^f$ | $(1 + \delta)w_{X_d X}$ | $\frac{2c(1+\delta)}{1+c}w_{X_d X}$ | $c(1 + \delta)w_{X_d X}$ |
| drive resident | $\lambda_f = 1$ | $\lambda_f = 2$ | $\lambda_f \text{ large}$ |
| $w_{mat}^f$ | $\frac{w_{X_d X}}{w_{X_d X_d}}$ | $\frac{w_{X_d X}}{w_{X_d X_d}}$ | $\frac{w_{X_d X}}{w_{X_d X_d}}$ |
| $w_{mat}^m$ | $\frac{1}{w_{X_d Y}}$ | $\frac{1}{w_{X_d Y}}$ | $\frac{1}{w_{X_d Y}}$ |
| $w_{pat}^f$ | $\frac{1}{1+\delta} \frac{w_{X_d X}}{w_{X_d X_d}}$ | $\frac{2}{(1+c)(1+\delta)} \frac{w_{X_d X}}{w_{X_d X_d}}$ | $\frac{1}{c(1+\delta)} \frac{w_{X_d X}}{w_{X_d X_d}}$ |

The two parts of inequality Eq(5) reflect the two pathways via which a rare X chromosome can increase in frequency in females, which are equally weighted (Figure 1). First, X chromosomes can be inherited from mother to daughter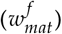. Second, X chromosomes in males are always inherited from the mother and will always then be passed to a daughter 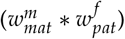. If, averaged over these two pathways, the frequency of female carriers increases, then a rare chromosome type will spread in the population.

**Figure 1.**
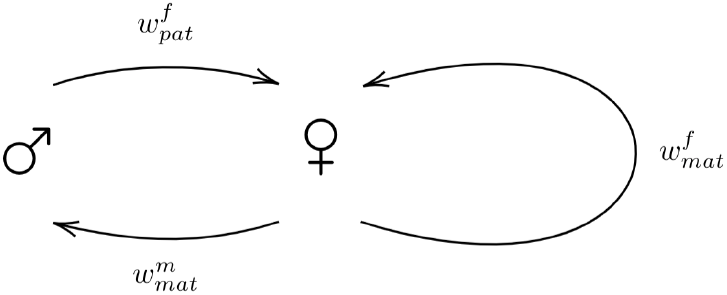
The paths a rare X chromosome variant can take in order to return to a female. All paternally inherited X chromosomes from an XY male will pass into females in the next generation. Females can transmit X chromosomes (maternally) to either sons or daughters. For a rare X chromosome type to spread in a population, it must increase in frequency in females, which may occur via either of the paths shown.

### Singly mating population

When females mate once (*λ_f_* = 1), a driving X_*d*_ chromosome will spread if

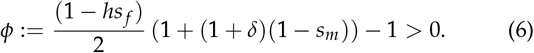

That is, X chromosome meiotic drivers spread if their their transmission advantage in males (1 + *δ*) is sufficiently large relative to their fitness cost in males and in female heterozygotes (*s_m_* and *hs_f_*). The driving X_*d*_ chromosomes will not fix in the population if

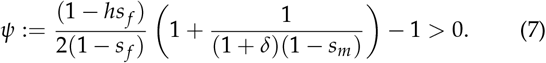

As X chromosome meiotic drive (X_*d*_) becomes common, the transmission and fitness advantage/disadvantage of X_*d*_ chromosomes in males is unchanged (terms involving *δ* and *s_m_*). Thus, the maintenance of polymorphism (satisfying inequalities in both Eq(6, 7)) occurs when meiotic drive causes low fitness cost in female heterozygotes (1 − *hs_f_*) relative to the cost in female homozygotes (1 − *s_f_*), which allows invasion but prevents fixation. For example, meiotic drive alleles are less likely to reach fixation when the negative fitness effects of drive are recessive (*h* = 0, Figure 2).

**Figure 2.**
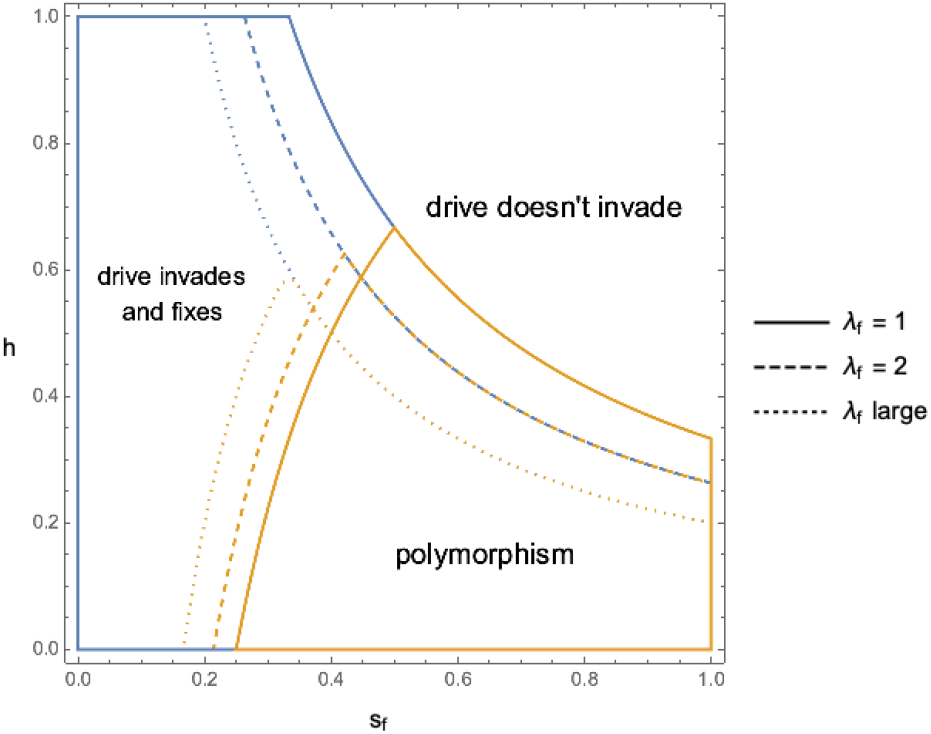
Fitness parameters under which X chromosome meiotic drive invades, reaches a polymorphism or fixes, for different levels of female selection and polyandry. Meiotic drive invades when heterozygous female carriers have sufficiently low fitness cost (*hs_f_* low). Meiotic drive will then also fix in the population unless the fitness cost in homozygotes (*s_f_*) is high. The level of polyandry is shown for one mating (*λ_f_* = 1, solid line), two matings (*λ_f_* = 2, dashed line), and multiple mating (*λ_f_* large, dotted line) where females effectively sample at random from all male sperm produced. Assuming that drive males produce less total sperm than non-drive males (*c* < 1), meiotic drive alleles invade and fix for a smaller parameter space when females are more polyandrous. Here *c* = 0.75, meiotic drive alleles have no fitness effects in males (*s_m_* = 0), and drive males produce only X_*d*_-bearing sperm (*δ* = 1), so the total fertility of drive males is reduced by an intermediate amount.

### Sperm competition in polyandrous populations

When females mate only once, the fertilization success of X chromosomes depends only on the ratio of X- to Y-bearing sperm, given by *δ* (Table 1). However, when females mate more than once, there is competition among sperm from multiple males. In this case, the fertility of drive males (*c*) is also important. When females mate twice (*λ_f_* = 2), the conditions for invasion and polymorphism (i.e. both X and X_*d*_ chromosomes) to be maintained by selection become

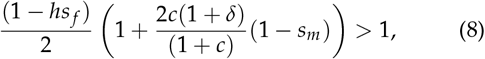

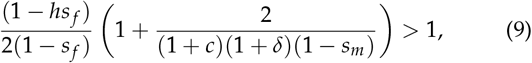

where (1 + *c*)/2 is the average quantity of sperm contributed by each male when a female mates with one drive male and one non-drive male. The conditions depend on drive male fertility *c*. The LHS of Eq(8) increases with *c*, as sperm compensation by drive males facilitates invasion by the driving X. However, the LHS of Eq(9) decreases with *c*, as compensation also aids fixation by meiotic drive. The net effect of compensation on the maintenance of polymorphisms can swing one way or another.

When females mate many times (*λ_f_* large), and effectively sample sperm at random from all males in the population, the conditions for X_*d*_ invasion and polymorphism respectively are

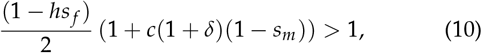

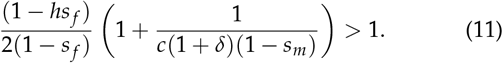

By comparing these conditions to inequalities Eq(8,9), we can see how drive male fertility (*c*) affects the propensity for a driving X to be polymorphic. Increased polyandry makes invasion harder because 2*c*/(1 + *c*) ≥ *c*. But polyandry promotes polymorphism given that a driving X is able to invade (1/*c* ≥ 2/(1 + *c*)).

When male carriers of meiotic drive produce the same overall quantity of sperm (full compensation, *c* = 1), their relative fertilization success is independent of sperm competition. The number of female mating partners doesn’t impact the invasion or fixation conditions for drive alleles (Eq(6, 8, 10) and Eq(7, 9, 11) are equivalent when *c* = 1). Without full compensation (*c* < 1), increasing female polyandry tends to disfavour both invasion and fixation of meiotic drive alleles as it increases sperm competition (Figure 2). With no compensation (sperm killers, *c* = 1/(1 + *δ*)), male drive-carriers produce the same quantity of X-bearing sperm as non-drive males. These sperm killing meiotic drive alleles are strongly disadvantaged by polyandry and cannot spread when females mate many times (inequality Eq(11) never satisfied).

### Limiting male matings narrows the polymorphism space

In the results presented above, we assumed that there is no sperm limitation, so even a small number of males is capable of fertilizing a large female population. In this case, extinction by meiotic drive only occurs when there are no males left in the population.

Here, we use the simulation model to consider limitations on the number of matings that a male can perform, which may limit population growth once the sex ratio becomes highly female biased. Figure 3 illustrates three different outcomes of the model on the assumption that individual males have a limit of 20 matings (*λ_m_* = 20). Parameter values are chosen under which a wild-type population is stably maintained (Figure 3A). However, when a driving X allele is introduced into the population it rapidly increases in frequency, which can skew the sex ratio further and further towards females until extinction ensues (Figure 3B). When the fitness costs of drive are higher, drive can be stably maintained. The resulting population is female-biased and larger than it would be in the absence of drive because the higher proportion of females increases the productivity of the population (Figure 3C).

**Figure 3.**
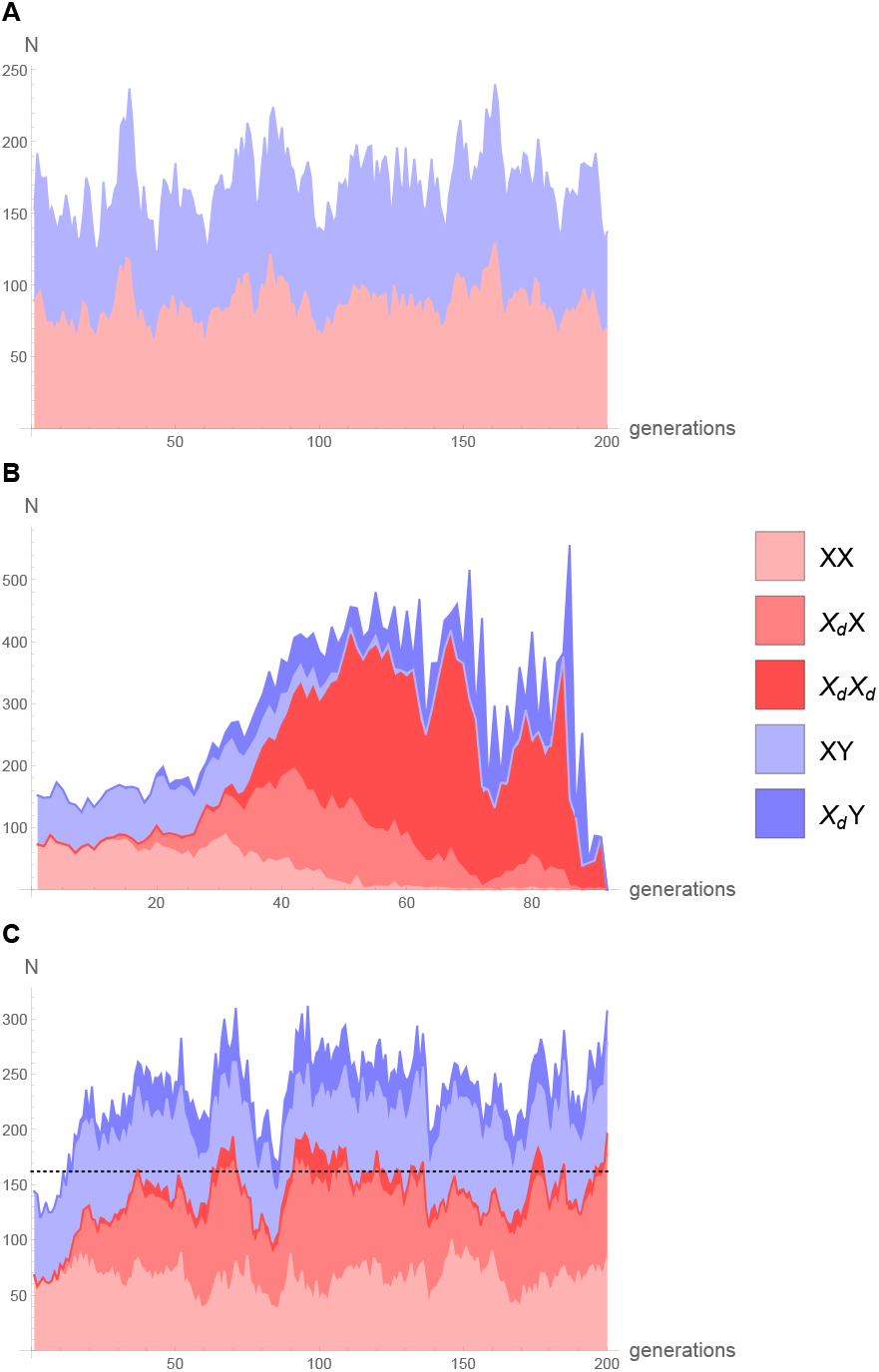
Illustrative examples of population dynamics with and without drive. a) the wild-type population without drive, b) the addition of drive causing rapid population extinction (*h* = 0.4, *s_f_* = 0.2), c) the addition of drive subject to stronger counter selection leading to a population polymorphic for drive (*h* = 0.2, *s_f_* = 0.55). The mean wild-type population size was 161 and is shown by the dotted line. Other parameters used were *c* = 0.75, *δ* = 1, *b* = 2.4, *α* = 10^−3^, *λ_f_* = 2, *λ_m_* = 20, *q* = 0.01, and the initial population size was 150.

The effect of limited male mating can be assessed by comparing the proportion of numerical simulations that result in drive polymorphism to the predictions from the analytical model, where there are no limits to male mating (Figure 4A). With male mating set at *λ_m_* = 20, the region of polymorphism shrinks. On the upper boundary, this represents conditions where the polymorphism is unstable because meiotic drive alleles have only a slight advantage and remain at low frequencies where they are exposed to loss by genetic drift. The leftmost boundary is where drive is strong enough to reach a high frequency. The sex ratio is heavily female biased, so many females go unmated due to male mating limitation, and the population can go extinct. This is illustrated in Figure 4B, where the maximum number of matings per male was reduced to *λ_m_* = 2. More populations go extinct close to the fixation boundary, but just as many lose drive stochastically.

**Figure 4.**
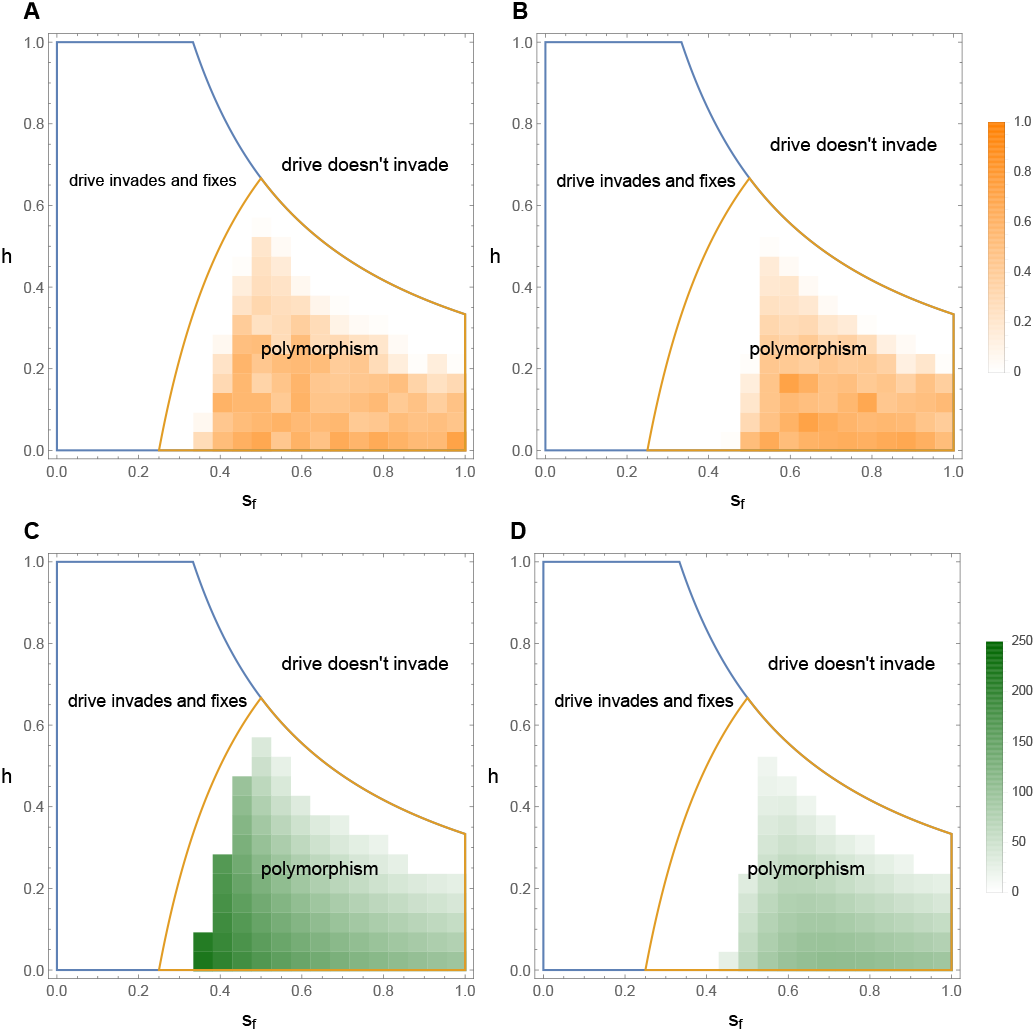
The effect of constraints on male mating rate. The region of polymorphism is demarcated on the assumption that there are no constraints on male mating (area within orange line). This is compared to numerical simulation data showing the proportion of times out of 50 populations that a polymorphism was maintained for 2000 generations when a) males mate 20 times (*λ_m_* = 20) and b) males mate twice (*λ_m_* = 2). The simulation parameters used were *δ* = 1, *c* = 1, *λ_f_* = 1, *q* = 0.01, *N*_0_ = 200, *b* = 2.4, *α* = 0.001. Figures c) and d) show the average increase in population size compared to a wild-type population without meiotic drive, over 2000 generations. The population size for each simulation was taken to be the mean size after a 100 generation burn in period, and the value for each tile in the plot is the mean of all simulations that resulted in polymorphism.

### Population size in the presence of drive

In this section we focus on the case where females mate only once, excluding the effects of sperm competition. In the absence of meiotic drive (*p* = 0), the population reaches an equilibrium population size 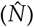 given by the intrinsic birth rate (*b*) and the density-dependent reduction in female fertility caused by competition among individuals (*α*):

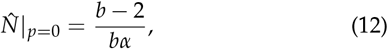

which is a standard result for logistic population growth with non-overlapping generations (Edelstein-Keshet 1987, pp.44-46 with *r* = *b*/2 and *d* = *bα*). The equilibrium population size is larger when the intrinsic birth rate (*b*) is higher or the competitive effect of other individuals (*α*) is weaker. For the population to persist, each female must produce at least two offspring (*b_min|p_*_=0_ = 2).

If an X chromosome meiotic driver invades (i.e. *ϕ* > 0, Eq(6)) and reaches a polymorphic equilibrium (i.e. *ψ* > 0, Eq(7)) then its frequency in females and males is given by

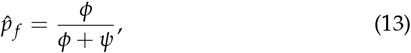

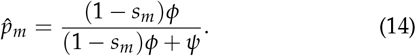

At the polymorphic equilibrium, the sex ratio will be female-biased and this in turn affects the ecological equilibrium population size

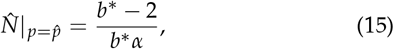

where

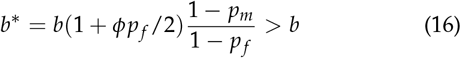

because *ϕ* and *p_f_* are non-negative and *p_m_* ≤ *p_f_* (from Eq(14)). As *b*^*^>*b*, the population size with drive is always larger than it would have been without drive (Figure 5). A similar outcome holds when a drive allele fixes, the total population size is then

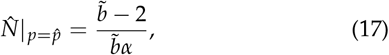

where

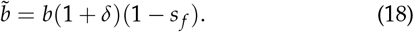

**Figure 5.**
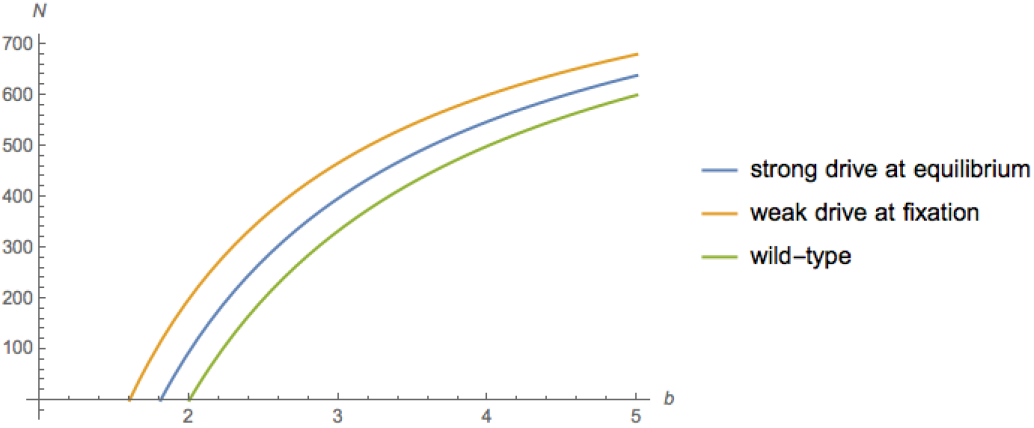
Population size with and without meiotic drive alleles given different birth rates (*b*). With meiotic drive, the population size is higher and can persist with lower intrinsic growth rates (*b* < 2), whether drive is weak (*δ* = 0.25) or strong (*δ* = 1). There were no drive costs to males, *s_m_* = 0, and males fully compensated for lost sperm (*c* = 1) so that female mating rate has no effect. The fitness costs associated with strong drive were *h* = 0.1, *s_f_* = 0.8, weak drive had no associated fitness cost (*s_f_* = 0). The density dependence was defined by *α* = 10^−3^.

For drive alleles that reach fixation, 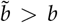. Again, by biasing the sex ratio towards females, fixed drive increases the population birth rate and thereby increases the overall population size (Figure 5). However, this result may be most relevant for weak meiotic drivers (*δ* < 1) because there will be no males in the population when strong meiotic drivers (*δ*≈1) reach fixation.

By increasing population productivity, meiotic drive alleles also help to protect populations from extinction. With strong drive at an intermediate equilibrium frequency, the minimum intrinsic birth rate required for population persistence is 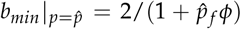, while for weak drive at fixation this is 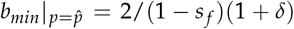. Both of these values are less than two, the cut-off value for a population to go extinct in the absence of drive (i.e. *b* < 2). Populations with drive can persist with a lower average number of offspring per female than those without, because a higher proportion of the population are female. The results of the simulation model align with the analytic model. Whenever a polymorphism is reached, the resulting population size is bigger than in the absence of drive (Figures 4C and D). The extent of the boost in population size depends on the viability cost associated with drive. As the cost decreases (either *h* or *s_f_* decreases), the equilibrium frequency of drive increases, the sex ratio becomes more female biased, and the increase in population size becomes larger.

Overall, these simulations confirm that meiotic drive can boost population sizes even when males can only fertilize a limited number of females. However, when meiotic drive causes the sex ratio to exceed the number of females that males can mate with, the presence of unmated females means population size declines and extinction risk increases.

### Population persistence time

Populations that are relatively small are liable to go extinct within a reasonable time due to demographic stochasticity. To examine the effect of drive on persistence times simulations were run in small populations with a low intrinsic birth rate (*b* = 2.4, *α* = 10^−2.4^), reflecting for example a small patch in a suboptimal or marginal environment. In these simulations, the mean population size without meiotic drive was 36.3 (SD: 12.7, the expected population size from Eq(12) was 41.9) and the mean persistence time was 1088 generations (coefficient of variation, *c_v_*: 0.92, (Everitt 1998)). The approximate alignment of the mean and standard deviation (ie, *c_v_*≈1) is expected because the persistence times of stochastic logistic growth models are exponential in distribution, and so the mean and standard deviation are approximately equal (Ovaskainen and Meerson 2010).

First, we consider the case where meiotic drive has no fitness costs (*s_f_* = *s_m_* = 0) and should either spread to fixation or be lost by drift (Figure 6A). With *δ* = 0 (i.e. no transmission distortion), the X_*d*_ allele is completely neutral and the population persists as if it were wild-type (Figure 6A). For increasingly strong meiotic drivers (increasing *δ*), the probability of invasion increases, and meiotic drive alleles are present at the end of a larger proportion of the simulations. Populations with drive persist for much longer both with weak and strong drive (0.05 < *δ* < 0.8) and the level of drive does not cause them to show an increase of the extinction rate (Figure 6A). However, when drive is very strong (*δ*≥0.8), the sex ratio can become excessively female biased and population extinction becomes more likely.

**Figure 6.**
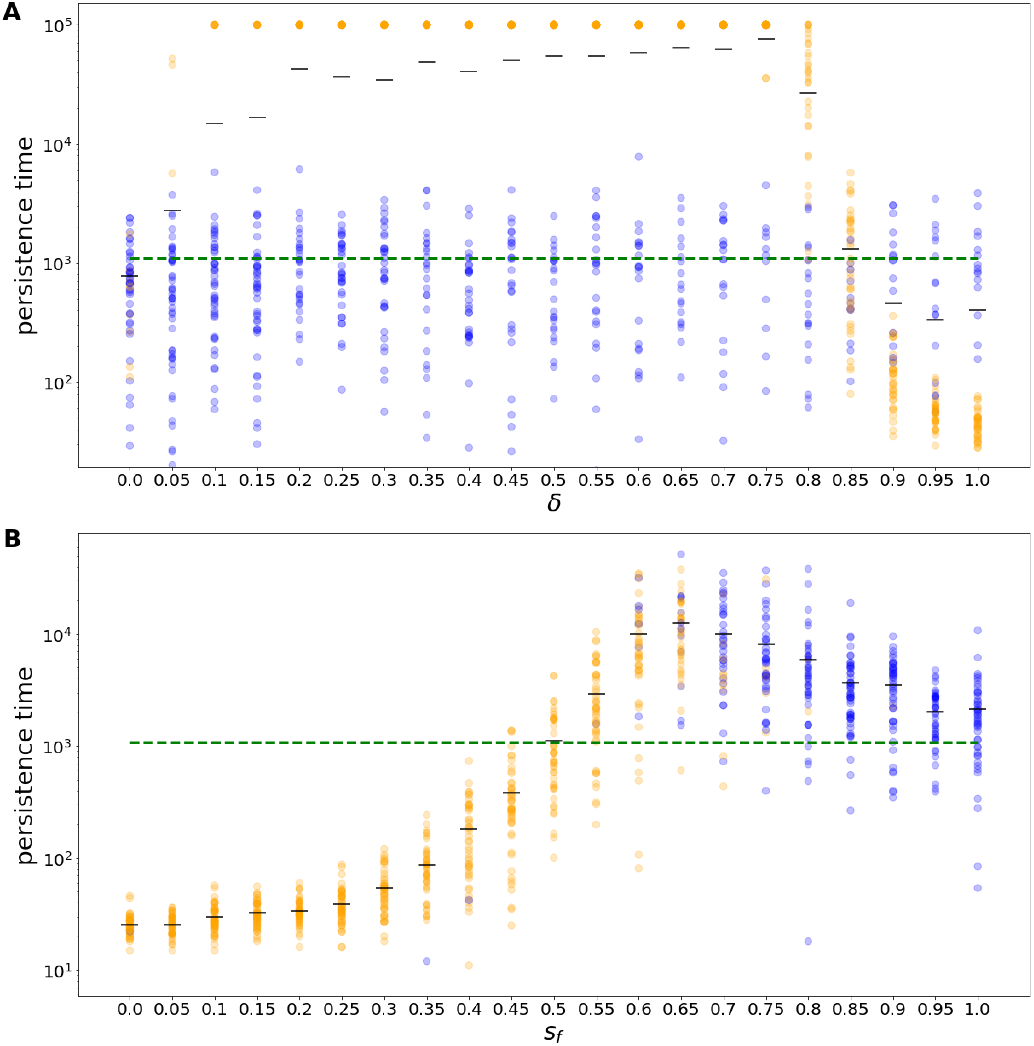
Persistence times for populations as (a) the strength of drive increases (*δ*), and (b) the strength of selection in females increases (*s_f_*). Orange points denote populations where drive was present and blue points where drive was absent at the time of extinction or at the maximum simulation duration of 10^5^ generations. The green dashed line in represents the mean persistence time of wild-type populations without meiotic drive and the black lines show mean persistence times. Populations began with an initial drive frequency of *q* = 0.1. Female adults had a mean birth rate of *b* = 2.4 with a high cost of competition, *α* = 10^−2.4^. In (a) *s_f_* = 0, drive acts by killing a fraction of Y sperm with no compensation (*c* = 1/(1 + *δ*)) and in (b) viability costs were in homozygotes only (*h* = 0), males produced only X_*d*_ sperm and had full compensation (*δ* = *c* = 1). Other parameter values *s_m_* = 0, *λ_f_* = 2, *λ_m_* = 20.

Population persistence was also evaluated for strong mei otic drivers (*δ* = 1). For simplicity, the dominance coefficient in females was set to *h* = 0, limiting viability reduction to homozygous female carriers (Figure 6B). When drive incurs no or small fitness costs (*s_f_* < 0.2), it spreads to fixation and causes rapid extinction through extreme sex ratios. As the cost increases (0.2 < *s_f_* < 0.5), meiotic drive reaches fixation more slowly and the sex ratio is less female biased. The persistence time increases back towards that found in wild-type populations. Even when drive is maintained at a polymorphic equilibrium, the frequency of drive can be high, giving heavily female biased sex ratios, which often results in extinction due to stochastic fluctuations in the sex ratio and reduces population persistence time. As the cost rises further (*s_f_* > 0.5), the frequency of drive at the polymorphic equilibrium falls and the sex ratio is less extreme, leading to longer population persistence than wild-type populations. Where the cost is very high (*s_f_* > 0.7), drive is maintained at a low equilibrium frequency and may itself be stochastically lost. However, the transient presence of drive still increases the overall longevity of the population.

## Discussion

A central finding of our analyses is that X-linked meiotic drivers generally increase population size. By biasing the sex ratio towards females, meiotic drive effectively boosts the population birth rate (Eq(16,18)). This increases the expected population size beyond the level in wild-type populations (Figures 4 and 5). In small populations at risk of stochastic population extinction, the increase in population size through meiotic drive can dramatically increase population persistence time (Figure 6). This should enable populations to persist in marginal environments where they would otherwise go extinct.

Female-biased sex ratios boost population size and persistence because birth rates are assumed to be limited by females. Thus, the population-level benefits of drive break down when males become scarce and are no longer able to mate often enough for females to achieve full fertility. Beyond this high level of sex ratio bias, the productivity of females falls, eventually below the population growth rate achieved in the absence of drive, and then to a point that threatens extinction (Figure 3B, Figure 6). Thus, to boost population size, strong meiotic drive must be held at a low frequency by countervailing selection, whereas weak meiotic drive can reach fixation and still increase population size and persistence (Figures 5, 6).

We derive general conditions for the invasion and fixation of X-linked alleles, taking account of tranmission bias via meiotic drive and/or sperm competition. Invasion by new driving X alleles (*ϕ*, Eq(6)) is a balance between the strength of meiotic drive (*δ*) and the viability costs in male carriers (1 − *s_m_*) and heterozygous females (1 − *hs_f_*). Whether or not drive alleles fix or remain polymorphic (*ψ*, Eq(7)) depends in addition on the viability of homozygous females (1 − *s_f_*).

These results apply in the absence of either polyandry or male fertility costs to drive (*c* = 1). When drive males have reduced fertility, due to a reduction in the number of sperm produced per ejaculate or related reasons (*c* < 1), increased polyandry (*λ_f_*) impedes both invasion (Eq(8,10) and fixation of drive (Eq(9,11), Figure 2), in line with previous results that suggest polyandry hinders the spread and fixation of meiotic drive alleles (Price *et al.* 2010, 2014; Holman *et al.* 2015).

Although there are few empirically obtained estimates for the fitness costs of X-linked drive, many of them are compatible with polymorphism according to our model. Female viability costs in *Drosophila* are often recessive but strong (*h* = 0 − 0.11, *s_f_* = 0.56 − 1, see Table 1 in (Unckless and Clark 2014) and (Larner *et al.* 2019; Dyer and Hall 2019)). A counterfactual is the estimate from the stalk-eyed fly *Teleopsis dalmanni* (Finnegan *et al.* 2019a) which found additivity and weaker viability loss in egg-to-adult viability, though the range on the dominance estimate is large. A limitation of attempts to measure fitness is that they are based on laboratory conditions that may distort the pressures that exist in natural populations. They also typically measure one component of fitness, for example survival over a particular life stage, neglecting others such as reproductive success. Furthermore, we note that these empirical estimates may be biased towards systems with strong meiotic drive (*δ*≈1) because weak meiotic drivers are less easy to detect (Burt and Trivers 2006).

Population persistence is predicted to increase exponentially with size (Ovaskainen and Meerson 2010) such that increases in population size caused by meiotic drive can cause a large increase in population persistence time (Figure 6). Therefore, we predict that populations with meiotic drive are more likely to be observed in marginal habitats where wild-type populations may go extinct. In natural populations, tests of this prediction may be confounded by a range of other factors associated with marginal habitats. For instance the rate of polyandry is likely to be lower in low quality environments and this will favour the spread of drive Pinzone and Dyer (2013); Finnegan (2020). A viable first experimental step may be to use lab populations to evaluate whether X-linked meiotic drive can increase population birth rates and/or rescue declining populations from extinction.

A relationship between sex ratios and population size/persistence is also not yet clearly established in species with temperature-dependent sex determination, despite similar predictions (Boyle *et al.* 2014; Hays *et al.* 2017). As predicted previously (Hamilton 1967), severely male limited populations should be quickly driven to extinction, which can occur in lab populations (Price *et al.* 2010) and may have been observed in a natural population (Pinzone and Dyer 2013). However, high male mating rates can facilitate population persistence in the face of extremely biased sex ratios. *Wolbachia* infection in butterflies can result in a sex ratio of 100 females per male, but these populations may persist because males can mate more than 50 times in a lifetime (Dyson and Hurst 2004).

The population dynamics of sex ratio distorting elements are thought to be influenced by their propensity to colonise new patches and drive them to extinction, i.e., metapopulation dynamics (Hatcher 2000). When drive is strong and confers little fitness cost in females, new populations cannot be established by drive genotypes because of the deficit in the numbers of males and resulting weak population growth. This could lead to cycling dynamics where colonisation by non-drive genotypes is needed to establish populations, which can then be invaded by drive genotypes whose spread is followed by extinction (Taylor and Jaenike 2003). These population level costs can decrease the overall frequency of selfish genetic elements across the metapopulation (Boven and Weissing 1999). Our results emphasise the potential for X-linked meiotic drivers to boost population sizes and persistence times, which we expect would increase the proportion of patches expected to have drive. It has also been suggested that individuals carrying selfish genetic elements may show a greater propensity to migrate between populations, increasing their fitness by reaching patches with lower numbers of heterozygotes and less polyandry (Runge and Lindholm 2018). However, the full metapopulation dynamics where local population sizes are affected by drive frequency remains to be investigated.

We generally predict population size to be increased when the sex ratio is biased towards females. Thus we expect our results to hold in species with ZW sex determination when meiotic drive favours W chromosomes (Kern *et al.* 2015) but not when meiotic drivers favours Y chromosomes or Z chromosomes (Hickey and Craig 1966; Gileva 1987). A general constraint on our conclusions is that they hold for competition models where an increase in birth rate increases population size (Supplementary Information). If the population is limited by the availability of resources regardless of the birth rate, boosts in population size are not expected. Likewise, where males contribute to parental care either through direct care or via control of resources used by females, sex ratio distortion will not have such a profound effect because the expected change in the number of offspring produced will be reduced and have a lesser effect on population size and persistence (West 2009).

Our results are also pertinent to the design of synthetic gene drive systems. Gene drive systems have been proposed as a method of controlling pest populations through altering the sex ratio so that one sex becomes limiting. Many of these proposals are analogous to Y-linked meiotic drive, for example “X-shredders” (Windbichler *et al.* 2008; Galizi *et al.* 2014; Burt and Deredec 2018) that limit the reproductive output of the population by biasing segregation towards Y-bearing sperm. We expect systems that cause male sex ratio bias to be effective. X-drive has also been recently suggested as a tool for biological control (Prowse *et al.* 2019). As observed in some simulations, as long as males are not limiting, the population may benefit from the introduction of an X-drive that increases the population productivity and carrying capacity (Prowse *et al.* 2019). That is, less efficient synthetic X-drivers may fix and result in larger populations without causing populations to crash (Prowse *et al.* 2019); this is analogous to fixation of weak meiotic drive in our model. Another possibility is that the driving allele does not fix but is maintained at a polymorphic equilibrium by the evolution of suppressors or associated fitness costs, for example. The resulting population will have a female-biased sex ratio, which our results suggest could increase population size and persistence. Thus, we urge caution when considering the use of X-linked gene drive for population control.

At the population level, the optimal sex ratio is likely to be female biased because relatively few males are required for complete fertilization. In some circumstances, such as local mate competition, individual-level and group-level selection can align, and female-biased sex ratios can evolve (West 2009; Hardy and Boulton 2019). Here, we show that selfish genetic elements (specifically, X-linked meiotic drivers) can move populations towards their population-level optimum and benefit population-level traits (such as population size and persistence time), a possibility that has probably been under-emphasised relative to their detrimental effects on populations.

## Acknowledgements

The authors would like to thank Max Reuter and Ewan Flintham for their comments on the manuscript. CM is supported by CoMPLEX and EPSRC studentship EP/N509577/1. AP is supported by Engineering and Physical Sciences Research Council grants (EP/F500351/1, EP/I017909/1), and Natural Environment Research Council grant (NE/R010579/1). MFS is supported by BBSRC grants BB/M011585/1 and BB/P024726/1.

## Author contributions

The research project was conceived, carried out and the paper written by all authors. CM and MFS carried out the modelling work.

## Supplementary Information

### Alternative form of density dependence

In the main text, we assumed that competition for resources among adults is a source of density dependent selection by reducing the fertility of adult females. Here, we explore an alternative form of density dependence in which competition for resources occurs among juveniles. In this model, *J* = *bN* is the number of juveniles produced in each generation. The survival probability for juveniles with genotype *i* is given by

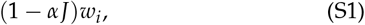

where *w_i_* is the viability fitness of the individual as described in Table 1 in the main text. The implementation of this fitness function is exactly the same as described in the main text but here we replace *N* with *J*.

**Figure S1.**
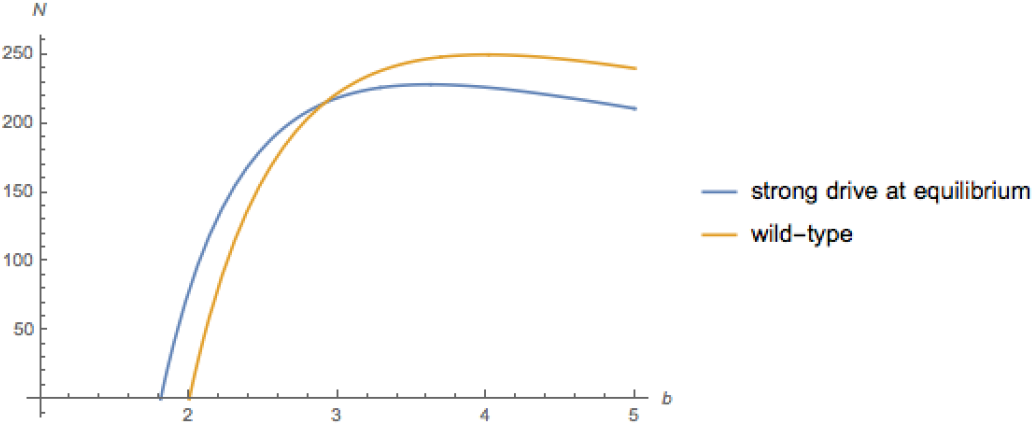
Population size with and without meiotic drive alleles given different birth rates (*b*) under an alternative form of density dependent selection. With meiotic drive, the population can persist with lower intrinsic growth rates (*b* < 2). There were no drive costs to males, *s_m_* = 0, and males fully compensated for lost sperm (*c* = 1) so that female mating rate has no effect. The fitness costs associated with strong drive were *h* = 0.1, *s_f_* = 0.8, and females were assumed to mate once, *λ_f_* = 1. The density dependence was defined by *α* = 10^−3^.

With resource competition among juveniles, increasing the birth rate does not always increase population size (Figure S1). Without meiotic drive, the equilibrium population size is

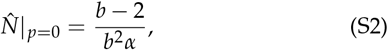

which includes a quadratic term in *b* not present when competition occurs among adults (Eq(12)). With juvenile competition, high birth rates both increase the number of juveniles, *J*, and increase the strength of competition among them. Thus, when birth rates are very high, the equilibrium population size decreases because competition among juveniles becomes intense.

As in our main results, we find that the intrinsic birth rate must be at least two for wild-type populations to persist whereas populations with drive can persist with a lower intrinsic birth rate (Figure S1). However, meiotic drive does not always increase population size in this scenario because increasing the effective birth rate by biasing the sex ratio towards females does not always lead to larger populations.

